# CD4^+^ T cell activation is dependent on a novel form of ULK1/2-independent autophagy

**DOI:** 10.64898/2026.04.29.721097

**Authors:** Edward Corrigan, Jan van der Beek, Brenda Raud, Julia Oliveira Lima, Ann de Mazière, Anna Knol, Cornelieke Pals, Derk Amsen, Judith Klumperman, Enric Mocholi, Paul J Coffer

## Abstract

Autophagy is essential for CD4+ T cell activation and immune regulation. However, during activation both autophagy and anabolic signaling must be simultaneously sustained, challenging established models of pathway antagonism. Here, we show that T cell receptor signaling and co-stimulation induce a non-canonical form of autophagy required for proliferation and cytokine production. Pharmacological and genetic analyses reveal that this pathway is activated concurrently with mTORC1, and is dependent on PIK3C3, but occurs independently of the canonical regulators ULK1/2, AMPK, ATG13, and Beclin 1. Furthermore, immuno-electron microscopy demonstrates that activation generates smaller autophagic structures that associate with multivesicular bodies and exhibit a unique morphology. These findings uncover a fundamental rewiring of autophagy control in CD4+ T cells and identify a novel form of mechanistically and morphologically distinct non-canonical autophagy.

## Introduction

Macroautophagy, hereinafter termed autophagy, is a highly conserved lysosome-mediated degradation pathway characterized by the encapsulation and trafficking of cellular components by double membrane vesicles called autophagosomes [1]. In CD4+ T cells it is essential in maintaining homeostasis, as well as playing a critical role in differentiation and activation [2,3]. CD4+ T cells regulate the adaptive immune response by directing immune cell activation and differentiation through cytokine production. For this, they must first be activated to proliferate, differentiate and perform their effector functions. Activation requires the stimulation and integration of T cell receptor (TCR), co-stimulation and cytokine signaling [4]. Ourselves and others have demonstrated that autophagy increases during T cell activation in response to TCR and co-stimulatory signaling (TCR/Co-stimulation), and is required for proper activation [5–10]. These studies have focused mainly on the responses of CD4+ T cells and their T_H_1 and T_H_2 effector subtypes. However, given the diversity of T cell subsets and their different effector functions and physiological responses, it is unclear if this applies to all subsets [11–13]. Recently, using a murine autophagy reporter model, it was described that upon TCR engagement CD8+ T cells suppress autophagy [14,15]. In addition, previous work has suggested that cytokine signaling also plays a role in regulating autophagy during activation of T_H_1 and T_H_2 CD4+ T cells [8]. More recently, however, using *in vitro* differentiated mouse cytotoxic T lymphocytes (CTLs), it was shown that interleukin 2 (IL-2) and IL-4 repress autophagy during activation while IL-7 and IL-15 help maintain it in naïve and memory cells [14]. These discrepancies in our understanding of the role and regulation of autophagy during activation between T cell subsets indicate that one model does not fit all, and suggest that autophagic responses could be more intricately regulated and diverse than previously understood.

Autophagy initiation is orchestrated by the unc-51-like autophagy-activating kinase 1/2 (ULK1/2) complex, which acts as the master regulator for autophagosome biogenesis and co-ordinates the localization of various pathway members [1]. ULK1/2 complex activity is regulated in a reciprocal signaling axis by the mechanistic target of rapamycin complex 1 (mTORC1) and the AMP-activated protein kinase (AMPK) complex; with mTORC1 inhibiting ULK1/2 under nutrient and growth factor-rich conditions, while AMPK activates it during energy stress [16–18]. A signaling paradox emerges during activation, whereby CD4+ T cells require high levels of mTORC1 activity and anabolism in order to expand, differentiate and produce cytokines [19], but also must simultaneously upregulate autophagy [8,9]. This suggests that an uncoupling in the AMPK-mTORC1-ULK1/2 regulatory axis could be occurring during activation. However, how CD4+ T cells sustain high autophagic flux despite elevated mTORC1-activity remains unknown.

Here, we present a distinct form of autophagy in CD4+ T cells that is both mechanistically and morphologically distinct, operating independently of the canonical AMPK-mTORC1-ULK1/2 axis. Understanding the regulatory mechanisms governing this pathway could not only provide critical insights into non-canonical autophagy but also enable the development of targeted therapeutic strategies to modulate CD4+ T cell activation.

## Results

### TCR/co-stimulation induces autophagic flux proportional to activation strength

To evaluate the effect of activation on autophagy, CD4+ T cells were isolated from the peripheral blood mononuclear cells (PBMCs) of healthy donors and stimulated with anti-CD3 and anti-CD28 to mimic TCR activation and co-stimulation, respectively. Lysosome-mediated autophagosome degradation rate (hereafter termed autophagic flux) was determined by the accumulation of microtubule-associated protein 1 light chain 3-II (LC3II) after treatment with bafilomycin A1 (BafA1) to block lysosome acidification. This was measured by either Western blot or flow cytometry with saponin extraction [20,21]. With both methods, a significant induction of autophagy was observed in CD4+ T cells after CD3/CD28 stimulation (**Figures 1A, 1B and S1A)**. The increase in autophagic flux in response to CD3/CD28 stimulation was dose dependent (**Figure 1C**), and proportional to IL-2 production (**Figure 1D**) and proliferation, as measured by Ki67 expression (**Figures 1E, S1B and S1C**) Similar autophagic responses were observed in both naïve (CD45RA+) and memory (CD45RO+) subsets (**Figures 1F, 1G and S1D**). However, both the basal and CD3/CD28-induced rate of autophagy were slightly higher in CD45RO+ cells (**Figure S1E and S1F**). Treatment with exogenous γ-chain cytokines IL-2, IL-7, or IL-15, had no effect on levels of autophagic flux in either unstimulated or activated cells (**Figures 1H-J and S2A-C**).

**Figure 1.**
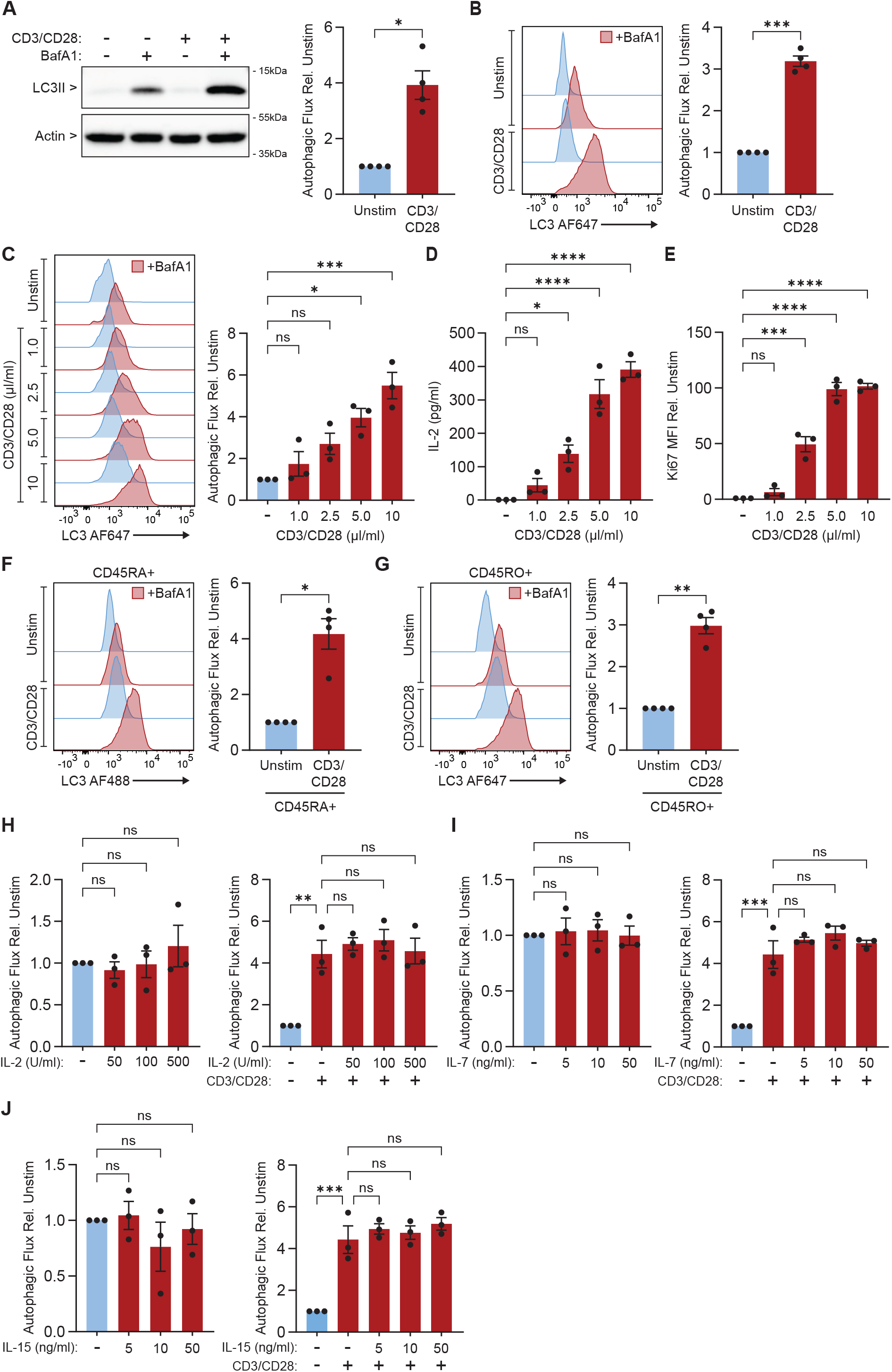
Autophagy is induced by TCR/co-stimulation, and unaffected by exogenous γ-chain cytokines in CD4+ T cells. (**A and B**) CD4+ T cells either stimulated 24 h with anti-CD3/CD28 or left unstimulated (Unstim) and treated with ± 100 nM bafilomycin A1 (BafA1) for 3 h. (**A**) Representative western blot of autophagic flux, as measured by LC3II turnover (n=4). (**B**) Representative normalized modal histograms of autophagic flux, measured through flow cytometry using saponin extraction and LC3II median fluorescence intensity (MFI) (n=4). (**C and D**) CD4+ T cells stimulated 24 h with increasing concentrations of anti-CD3/CD28. Panels show: (**C**) Representative normalized modal histograms of autophagic flux measured through flow cytometry using saponin extraction and LC3II MFI, cells were treated with ± 100 nM BafA1 for 3 h (n=3); and (**D**) IL-2 production measured by IL-2 ELISA (n=3). (**E**) Cell proliferation in CD4+ T cells stimulated 48 h with increasing concentrations of anti-CD3/CD28, determined by Ki67 MFI and measured through flow cytometry (n=3). (**F and G**) Representative normalized modal histograms of autophagic flux, measured through flow cytometry using saponin extraction and LC3II MFI, of CD45RA+ sorted CD4+ T cells and (**G**) CD45RO+ sorted CD4+ T cells from the same donor, stimulated 24 h with anti-CD3/CD28 and treated with ± 100 nM BafA1 for 3 h (n=4). (**H-J**) Quantifications of autophagic flux measured through flow cytometry using saponin extraction and measuring LC3II MFI of CD4+ T cells either left unstimulated or stimulated 24 h with anti-CD3/CD28, treated with ± 100 nM BafA1 for 3 h, and either (**H**) increasing concentrations of IL-2, (**I**) increasing concentrations of IL-7, or (**J**) increasing concentrations of IL-15 (n=3). Please note, for some cytokine treatments experiments were performed at the same time with the same donor and thus share the same control conditions. All graphs represent mean ± SEM. Statistical significance was measured by one-way ANOVA with Tukey post-hoc test, or by Welch’s t-test. ∗*p* < 0.05, ∗∗*p* < 0.01, ∗∗∗*p* < 0.001, ∗∗∗∗*p* < 0.0001.

Taken together, these results show that T cell receptor and co-stimulation signaling induces autophagy in naïve and memory CD4+ T cells in proportion to increases in IL-2 production and proliferation. While, exogenous γ-chain cytokines IL-2, IL-7, or IL-15 have no effect on levels of autophagy induction.

### TCR/co-stimulation-induced autophagy bypasses the canonical mTORC1-ULK1/2 inhibitory axis

During activation, CD4+ T cells require simultaneously high levels of autophagy and mTORC1-mediated protein synthesis [8,19]. Following CD3/CD28 stimulation, upregulation of both autophagic flux and mTORC1 activity, as measured by increased phospho-ribosomal protein S6 (S6) s235/236 levels, was observed (**Figure 2A**). To understand how these normally antagonistic pathways could be activated simultaneously, the AMPK-mTORC1-ULK1/2 signaling axis was interrogated (**Figure S3C**). In unstimulated cells, inhibition of mTOR with Rapamycin (RAP) and Torin 1 failed to increase autophagic flux (**Figure 2B**), suggesting a breakdown in the canonical ULK1/2-mTORC1 relationship [17]. Despite an increase in autophagic flux, CD3/CD28 stimulation resulted in increased phosphorylation of ULK1/2 at the mTORC1-dependent inhibitory site s757 [22] (**Figure 2C**). AMPK was also activated following CD3/CD28 stimulation, as indicated by increased pAMPK t172 and phospho-acetyl-CoA carboxylase (ACC) s79 phosphorylation levels (**Figures 2D and S3B**). To evaluate the potential role of AMPK, phosphorylation of ULK1/2 s555, an AMPK-target site associated with autophagy induction [22], was assessed. For these experiments, CD4+ T cells were additionally cultured in low fetal bovine serum (FBS) to induce amino acid starvation, under which autophagy is expected to be elevated and mTORC1 activity reduced. pULK1 s555 levels remained unchanged following either CD3/CD28 stimulation or FBS starvation however (**Figure 2E**), further supporting a breakdown in the canonical AMPK-ULK1/2-mTORC1 signaling axis.

**Figure 2.**
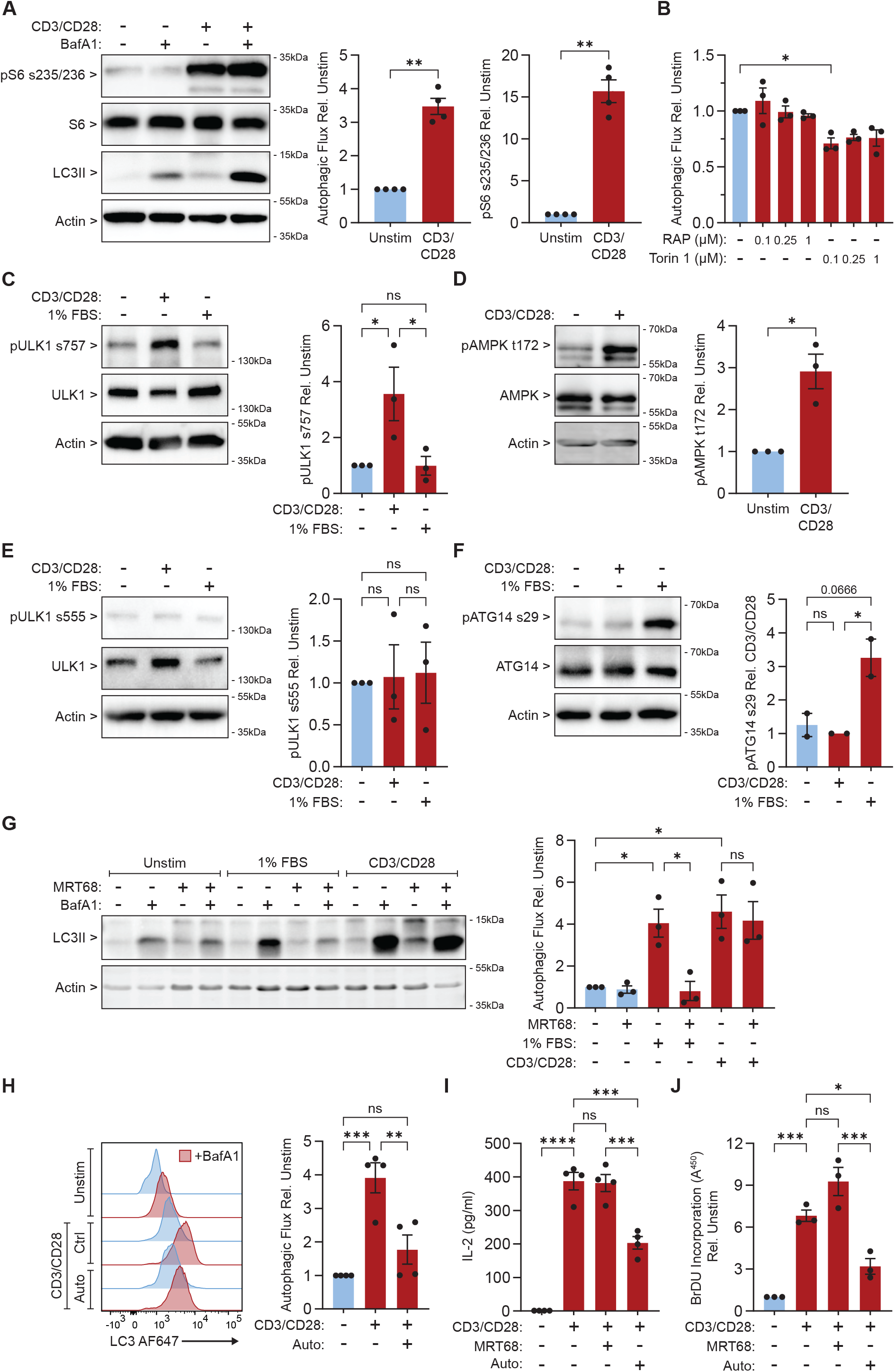
Autophagy induced by TCR/co-stimulation in CD4+ T cells is mTORC1 and ULK1/2-independent, but PIK3C3-dependent. (**A**) CD4+ T cells stimulated 24 h with anti-CD3/CD28 and treated with ± 100 nM BafA1 for 3 h. Representative western blot of autophagic flux and mTORC1 activity, as measured by LC3II turnover and pS6 s235/236 levels (n=4). (**B**) CD4+ T cells treated with increasing concentrations of Rapamycin (RAP) and Torin 1 for 24 h and treated with ± 100 nM BafA1 for 3 h. Autophagic flux measured by flow cytometry using saponin extraction and LC3II MFI (n=4). (**C-F**) CD4+ T cells stimulated with anti-CD3/CD28 or 1% FBS-starved 24 h. Representative western blots with quantifications of (**C**) pULK1 s757, (**D**) pAMPK t172 and (**E**) pULK1s555 (n=3), and (**F**) pATG14 s29 (n=2). (**G**) CD4+ T cells stimulated with anti-CD3/CD28 or 1% FBS starved and treated with ± 1 µM MRT68921 (MRT68) 24 h and ± 100 nM BafA1 for 3 h. Representative western blot of autophagic flux, with quantifications (n=3). (**H**) CD4+ T cells stimulated with anti-CD3/CD28 and treated with ± 1 µM Autophinib (Auto) 24 h and ± 100 nM BafA1 for 3 h. Representative normalized modal histograms of autophagic flux, measured through flow cytometry using saponin extraction and LC3II MFI (n=4). (**I and J**) CD4+ T cells stimulated 24 h with anti-CD3/CD28 and treated with 1 µM Auto or 1 µM MRT68. Graphs show: (**I**) IL-2 production measured by ELISA (n=4), and (**J**) proliferation measured by BrDU incorporation through colorimetric BrDU ELISA (n=3). All graphs represent mean ± SEM. Statistical significance was measured by one-way ANOVA with Tukey post-hoc test, or by Welch’s t-test. ∗*p* < 0.05, ∗∗*p* < 0.01, ∗∗∗*p* < 0.001, ∗∗∗∗*p* < 0.0001.

ULK1/2 regulates autophagy in part through the phosphorylation of autophagy-related gene 14 (ATG14) at s29 [23]. The promotes the assembly of the phosphatidylinositol 3-kinase catalytic subunit type 3 complex 1 (PIK3C3-C1), consisting of ATG14, Beclin 1, phosphoinositide-3-kinase regulatory subunit 4 (PIK3R4) and PIK3C3 (**Figure S3C**). In CD4^+^ T cells, ATG14 S29 phosphorylation increased during FBS starvation but not following CD3/CD28 stimulation (**Figure 2F**). To establish whether autophagy proceeds independently of ULK1/2, CD4+ T cells were either starved or stimulated with anti-CD3/CD28 in the presence of ULK1/2 inhibitor MRT68921 (MRT68). ULK1/2 inhibition blocked autophagy during FBS starvation, but not following CD3/CD28 stimulation, where autophagic flux still increased (**Figure 2G**). This was confirmed using an alternate ULK1/2 inhibitor, ULK101 (**Figure S3D**).

Collectively these findings demonstrate an uncoupling of the canonical AMPK-ULK1/2-mTORC1 signaling axis in CD4+ T cells. While starvation-induced autophagy is dependent on ULK1/2, during activation TCR/Co-stimulation engages a non-canonical pathway that is independent from ULK1/2 and mTORC1-mediated inhibition.

### TCR/co-stimulation-induced autophagy is dependent on PIK3C3

In the absence of ULK1/2 activity, we sought to identify which pathway components were still required for autophagy during CD4+ T cell activation. ULK1/2 initiates the formation of PIK3C3-C1, enhancing PIK3C3 lipid kinase activity and driving the production of phosphatidylinositol 3-phosphate (PI(3)P). PI(3)P subsequently facilitates autophagosomal membrane expansion through the recruitment of PI(3)P-binding proteins [24]. Previous work in mouse primary, and T_H_1 differentiated, CD4+ T cells has shown that inhibition of PIK3C3 can impair T cell activation [7,9,25]. To assess whether PIK3C3 is essential for TCR/co-stimulation-induced autophagy, cells were treated with the selective PIK3C3 inhibitor Autophinib (Auto). Autophagy induction by CD3/C28 was significantly inhibited by Auto (**Figure 2H**). This was confirmed using an alternate PIK3C3 inhibitor, PIKIII (**Figure S3E**). To evaluate whether CD4+ T cell activation is dependent on ULK1/2 or PIK3C3, IL-2 production, proliferation and expression of activation markers CD25 and CD69, was measured in cells stimulated with anti-CD3/CD28 in the presence of MRT68 or Auto. MRT68 had no significant effect on IL-2 production or proliferation, while Auto lead to a significant reduction in both (**Figures 2I and 2J**). This indicates that CD4+ T cell activation can occur independently of ULK1/2 but is contingent on PIK3C3-dependent autophagy. Neither ULK1/2 or PIK3C3 inhibition had a significant effect on activation-related surface marker expression (**Figures S3F, S3G and S3H**).

PIK3C3 is a class III phosphoinositide 3-kinase (PI3K) and part of the PI3K family, which includes class I PI3Ks that are essential for other T cell activation-related pathways such as the PI3K/AKT/mTORC1 (PAM) pathway [26]. To assess PIK3C3 inhibitor specificity, IL-7 was utilized as a control stimulus to activate PAM signaling, as it does not stimulate autophagy in our system (**Figure 1I**). PAM pathway activity was assessed by measuring phospho-AKT1 s473 and pS6 s235/236, with the broad spectrum PI3K inhibitor LY294002 (LY2) included as a positive control. Treatment with Auto of PIKIII had no significant effect on pAKT s473 or pS6 s235/236 levels following IL-7 stimulation, indicating the inhibitors had no off-target effects on PI3K signaling (**Figure S4A**).

To investigate whether this distinct autophagy program could also be induced by alternative activation methods, CD4+ T cells were stimulated with phorbol 12-myristate 13-acetate (PMA) and ionomycin (ION) which bypass the need for receptor binding by inducing downstream TCR/co-stimulation pathways directly to activate T cells [27,28]. As with anti-CD3/CD28 stimulation, PMA/ION treatment increased autophagic flux together with S6 s235/236 phosphorylation. This response was also ULK1/2-independent and PIK3C3-dependent (**Figures S4B and S4C**), indicating that activation of a subset of downstream signaling pathways alone is sufficient to drive this ULK1/2-independent non-canonical autophagy program.

These results reveal that TCR/co-stimulation-induced autophagy is PIK3C3-dependent, and demonstrate that it has an essential role for cytokine production and proliferation during CD4+ T cell activation.

### TCR/co-stimulation-induced autophagy is independent of AMPK, ATG13 and Beclin 1

To better understand how CD3/CD28-induced autophagy remains PIK3C3-dependent in the absence of ULK1/2 activity, we examined the role of key pathway members with regulatory potential. AMPK, which is activated by CD3/CD28 stimulation (**Figure 2C**), has been reported to regulate autophagy independently of ULK1/2 through direct and indirect interactions with Beclin 1 and PIK3C3 [29,30]. To determine whether CD3/CD28-induced autophagy is regulated by AMPK, CD4+ T cells were stimulated with anti-CD3/CD28 in the presence of AMPK inhibitor BAY-3827 (BAY). AMPK inhibition had no effect on autophagic flux following CD3/CD28 stimulation, indicating this pathway is AMPK-independent (**Figure 3A**). The efficacy of BAY to inhibit AMPK during CD4+ T cell activation was validated using ACC s79 phosphorylation as a marker of activity (**Figure S5A**). Consistent with these observations, AMPK inhibition did not affect FBS starvation-induced autophagy either (**Figure S5B**).

**Figure 3.**
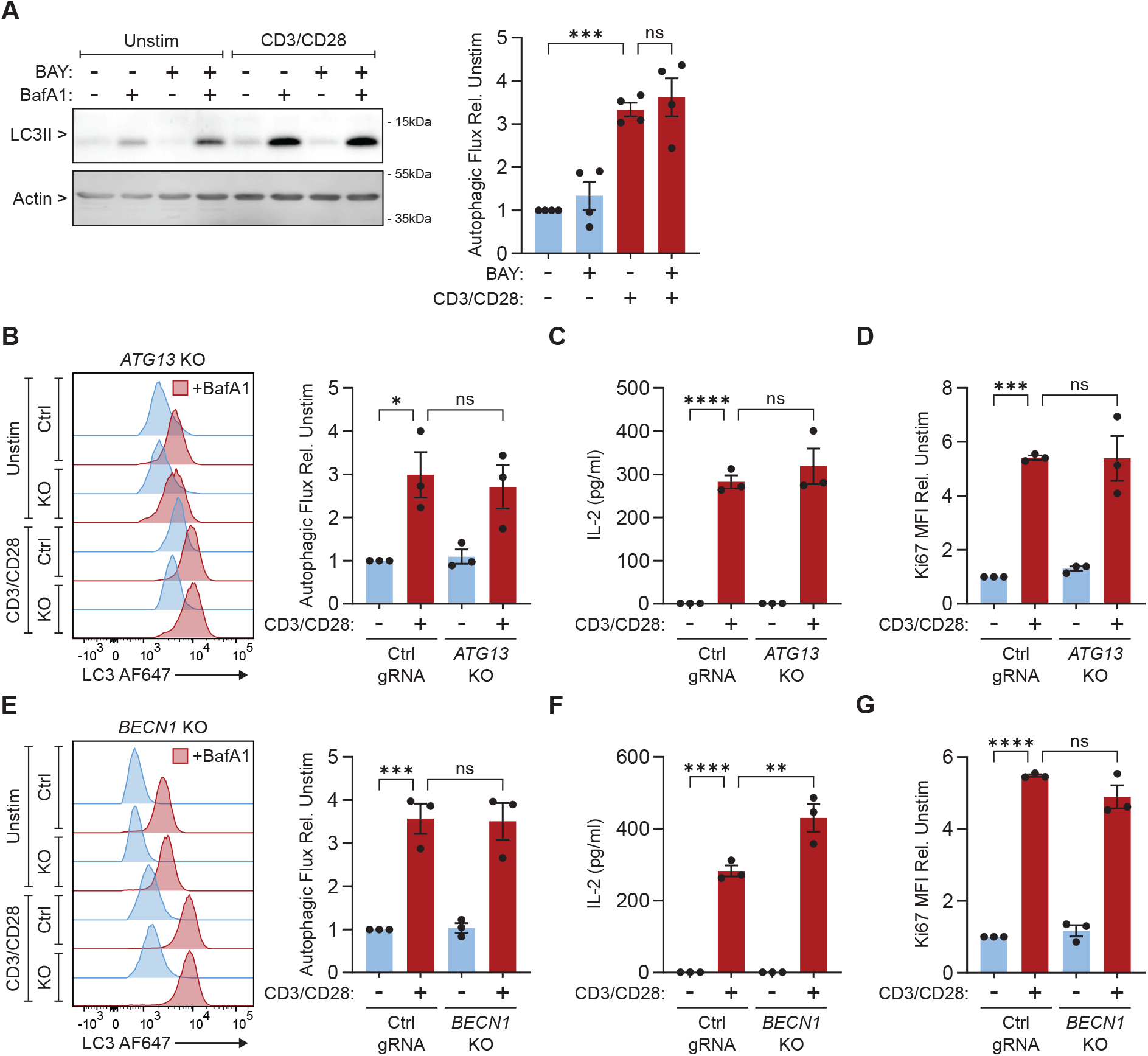
Autophagy induced by TCR/co-stimulation in CD4+ T cells is AMPK, ATG13 and Beclin 1-independent. (**A**) CD4+ T cells stimulated with anti-CD3/CD28 and treated with ± 0.5 µM BAY-3827 (BAY) 24 h and ± 100 nM BafA1 for 3 h. Representative western blot of autophagic flux, with quantifications (n=4). (**B-D**) Non-targeting control treated (Ctrl gRNA) and *ATG13* knockout (KO) CD4+ T cells stimulated with anti-CD3/CD28 24 h. Panels show: (**B**) Representative normalized modal histograms of autophagic flux measured through flow cytometry using saponin extraction and LC3II MFI, of cells treated ± 100 nM BafA1 for 3 h, with quantifications (n=3); (**C**) IL-2 production measured by ELISA (n=3); and (**D**) proliferation measured by Ki67 MFI using flow cytometry (n=3). (**E-G**) Ctrl gRNA and *BECN1* KO CD4+ T cells stimulated with anti-CD3/CD28 24 h. Panels show: (**E**) Representative normalized modal histograms of autophagic flux measured through flow cytometry using saponin extraction and LC3II MFI, of cells treated ± 100 nM BafA1 for 3 h, with quantifications (n=3); (**F**) IL-2 production measured by ELISA (n=3); and (**G**) proliferation measured by Ki67 MFI using flow cytometry (n=3). Please note, some *ATG13* KO and *BECN1* KO experiments were performed at the same time, with the same donor, and thus share the same Ctrl gRNA conditions. All graphs represent mean ± SEM. Statistical significance was measured by one-way ANOVA with Tukey post-hoc test, or by Welch’s t-test. ∗*p* < 0.05, ∗∗*p* < 0.01, ∗∗∗*p* < 0.001, ∗∗∗∗*p* < 0.0001.

The ULK1/2 complex is composed of ATG13, RB1-inducible coiled-coil protein 1 (FIP200) and ATG101 [1] (**Figure S3C**). ATG13 has previously been shown to interact with FIP200 and ATG101 independently of ULK1/2 during autophagy induction [31,32]. To assess whether this complex remains functional in the absence of ULK1/2 activity, CRISPR-mediated *ATG13* knockout (KO) human CD4+ T cells were generated (**Figure S5C**). Following anti-CD3/CD28 stimulation, *ATG13* KO had no significant effect on autophagic flux (**Figure 3B**), IL-2 production (**Figure 3C**) or proliferation (**Figure 3D and S5D**), indicating that ATG13 is not required for regulation of autophagic flux or CD4+ T cell activation.

Beclin 1 is a central scaffolding component of PIK3C3-C1, functioning as both a signaling hub and regulator of PIK3C3 lipid kinase activity [24,33]. It is primarily regulated by the ULK1/2-AMPK-mTORC1 signaling axis [34,35] but can also be regulated through alternative mechanisms, such as disruption of its interaction with B Cell Lymphoma 2 (BCL-2) [36,37]. To assess its role in TCR/co-stimulation-induced autophagy and understand to what extent PIK3C3-C1 complex formation is still required for autophagy induction, CRISPR-mediated *BECN1* KO human CD4+ T cells were generated (**Figure S5E**). Following CD3/CD28-stimulation, *BECN1* KO had no significant effect on autophagic flux (**Figure 3E**) but did increase IL-2 production (**Figure 3F**), while having no effect on proliferation (**Figure 3G and S5F**). These results demonstrate that *BECN1* KO is not required for TCR/co-stimulation-induced autophagy to occur.

Together these findings further support that autophagy is regulated by a mechanistically distinct pathway that is independent of Beclin 1, ATG13 and AMPK, further underscoring the breakdown of the canonical AMPK-ULK1/2-mTORC1 signaling axis in CD4+ T cells.

### TCR/co-stimulation-induced autophagy is morphologically distinct

To gain further insight into the mechanism and regulation of TCR/Co-stimulation-induced autophagy, we investigated the morphology of autophagic structures during activation. CD4+ T cells were labelled for LC3 with gold particles and visualized using immuno-electron microscopy (EM) on cryosections, which allowed for the unambiguous identification of autophagic compartments despite their ultrastructural heterogeneity. Cells were left unstimulated, FBS-starved, or stimulated with PMA/ION, and treated with BafA1 for the final 2 h prior to fixation and processing for LC3 immuno-EM as previously described [42]. In both unstimulated and FBS-starved cells, LC3-positive structures were identified representing autophagosomes and autolysosomes, containing diverse cargo, including organelles (**Figure 4A, white arrowhead**) and cytosolic material (**Figures 4A and 4B**), which remained undegraded in the presence of BafA1. In contrast, PMA/ION-stimulated cells exhibited clear differences in the morphology and cargo composition of autophagic compartments (AC), with their membranes being more densely labelled with LC3 and their contents being generally more electron-dense and reminiscent of condensed cytosolic material rather than organelles (**Figure 4C**). Furthermore, densely labelled LC3+ material was also observed more in multivesicular bodies following PMA/ION-stimulation, which likely represented amphisomes forming from the fusion of autophagosomes and late endosomes [38] (**Figure 4C**). Quantification of LC3+ compartments revealed that PMA/ION-induced autophagic structures were significantly smaller than those formed after FBS starvation (**Figure 4D**), and that the overall LC3-labelling count per cell was significantly higher in activated cells (**Figure 4E**). As previously described non-canonical autophagy pathways have been characterized by the conjugation of LC3 to single membranes (CASM) [39], LC3+ membrane architecture was also evaluated. However, we found no evidence to suggest the LC3-positive structures represented CASM (**Figures 4A, 4B and 4C**).

**Figure 4.**
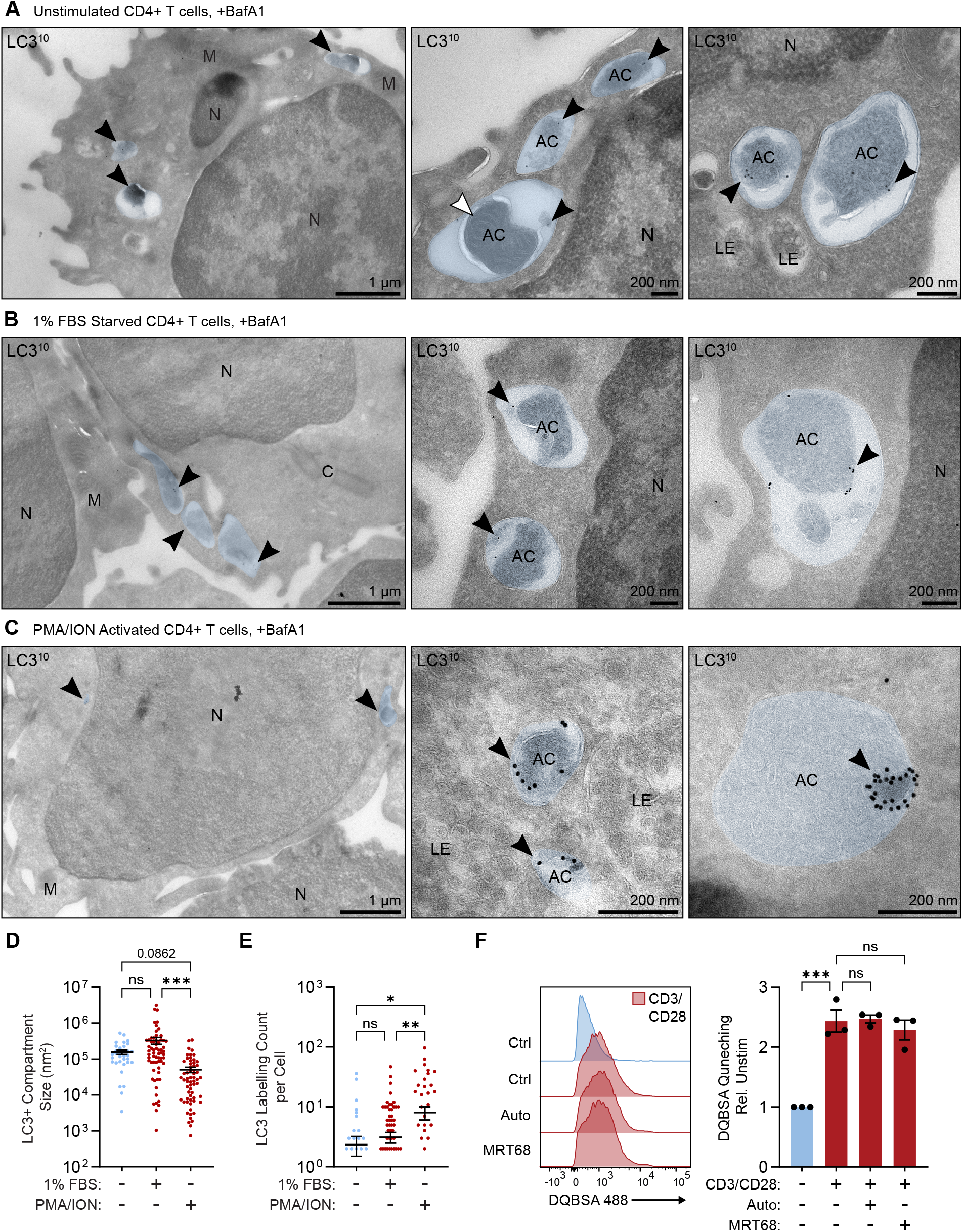
TCR/co-stimulation induces morphologically distinct autophagy, and PIK3C3-independent endolysosomal activity in CD4+ T cells. (**A-C**) Representative immuno-electron microscopy (EM) images labelled with LC3 and protein A 10 nm gold particles (PAG10) of CD4+ T cells either (**A**) unstimulated, (**B**) starved with 1% FBS, or (**C**) activated with PMA/ION, for 4 h and treated with 20 nM BafA1 for 2h prior to fixation. Black arrowheads denote LC3 gold positive membranes, blue coloring highlights LC3 gold positive compartments, and white arrowheads denote compartments containing organelles. AC = Autophagic compartment; C = Centriole; LE = Late endosome; M = Mitochondria; N = Nucleus. Scale bars: 1 µm or 200 nm where indicated. (**D and E**) Quantification of EM images (**A-C**), results are displayed on a log_10_ axes, bars indicate the mean value. (**D**) LC3 gold positive compartment size (nm^2^). (**E**) Membrane bound LC3 gold labelling count per cell. Due to log axis, 0 value labelling counts are not displayed on graph, but are still included in significance and mean value calculation. (**F**) Representative normalized modal histograms of DQ-BSA Green (DQBSA) dequenching, with n=3 DQBSA MFI quantifications, of CD4+ T cells stimulated 24 h with anti-CD3/CD28 and treated with 1 µM Auto or 1 µM MRT68, and cultured 6 h with 10 µg/ml DQBSA. All graphs represent mean ± SEM. Statistical significance was measured by one-way ANOVA with Tukey post-hoc test. ∗*p* < 0.05, ∗∗*p* < 0.01, ∗∗∗*p* < 0.001.

With the exception of PIK3C3, TCR/co-stimulation-induced autophagy has been shown to occur independently of all canonical regulatory components investigated. PIK3C3 has additional non-autophagy functions in the endolysosomal pathway, where it regulates endocytosis and vesicular trafficking [40–42]. Given the presence of LC3+ material within multivesicular bodies in PMA/ION-stimulated CD4+ T cells, we assessed the impact of PIK3C3 inhibition on endolysosomal activity to determine if the effects of PIK3C3 inhibition observed were due to endolysosomal inhibition. This was measured by flow cytometry using DQ-BSA Green (DQBSA) dequenching as a readout. MRT68 was included to evaluate the effect of ULK1/2 inhibition. CD3/CD28-stimulation significantly increased endolysosomal activity, as indicated by DQBSA-dequenching, while inhibition of PIK3C3 or ULK1/2 inhibition had no effect (**Figure 4F**). These findings were confirmed using alternative inhibitors PIKIII and ULK101 (**Figure S5G**). The DQBSA assay was validated using BafA1 to block lysosomal acidification and confirm lysosome-dependent dequenching (Figure S5H). These results reveal that TCR/co-stimulation-induced endolysosomal activation can occur independently of PIK3C3. This indicates that the effect of PIK3C3 inhibition on TCR/Co-stimulation-induced autophagy is not due to inhibition of PIK3C3-dependent endolysosomal activity, and is specific to the autophagy pathway.

Taken together, these results reveal a mechanistically and morphologically distinct form of autophagy occurs during CD4+ T cell activation, that is characterized by the formation of smaller autophagic structures that tend to form in multivesicular bodies. This pathway is dependent on PIK3C3, however this is not due to the role of PIK3C3 in the endolysosomal pathway, supporting the idea that a unique PIK3C3-dependent autophagy pathway is activated by TCR/Co-stimulation.

## Discussion

Here we describe a novel autophagy pathway induced in CD4+ T cells during activation by TCR/co-stimulation, that operates independently of mTORC1 and the major autophagy regulators AMPK and the ULK1/2 complex. These findings provide insight into how anabolic signaling and autophagy can occur simultaneously. Prior studies in CD4+ T cells and their effector subtypes have predominantly shown that autophagy increases in response to activation [7–9]. However discrepancies have emerged in the field with regards to the induction and level of autophagy in naïve and activated T cells, with differing autophagic responses being observed in CD8+ T cells [12–14]. These inconsistencies also extend to the role of cytokines in inducing and regulating autophagy during activation. Notably certain γ-chain cytokines have been shown to have opposing roles during activation between CD4+ and CD8+ T cell subsets [8,14]. Here we observed that both naïve and memory CD4+ T cells displayed low levels of autophagy when unstimulated, and that TCR/co-stimulation induced robust increases in autophagic flux in both subsets (**Figures 1F and 1G**). While their phenotypes were similar, memory cells had higher levels of basal autophagy and an increased response to CD3/CD28-stumulation (**Figure S1E and S1F**). This could be a result of memory cells having a higher resting metabolic rate, and a stronger and more rapid response to antigen stimulation [43]. We observed that exogenous IL-2, IL-7 and IL-15 had no effect on the levels of autophagy in CD4+ T cells (**Figures 1H, 1I and 1J**). This is consistent as in unstimulated cells, γ-chain cytokines activate the PAM signaling pathway (**Figure S4A**), stimulating negative autophagy regulators including mTORC1 [44–48]. While we demonstrate that mTORC1-independent autophagy can occur in CD4+ T cells (**Figures 2A and 2C**), the pathway described here requires the integration of TCR/co-stimulation pathways during activation and is not observed following γ-chain cytokine stimulation. The lack of cytokine-driven autophagy is in contrast to previous observations with *in vitro* differentiated mouse T_H_1 and T_H_2 effector CD4+ T cells [8]. This may indicate that context-dependent autophagic responses exist among CD4+ T cell effector subtypes or alternatively reflect species-specific differences or the effects of *in vitro* differentiation protocols that render cells more responsive to certain cytokine signals. Differences in autophagy between CD4+ and CD8+ T cell subsets would be plausible given they are discrete cell types, and that upon activation CD4+ T cells differentiate into distinct effector T cell subsets, whereas CD8+ T cells gain cytotoxic effector functions [49]. This could translate to distinct autophagic functions during activation; however, this still requires a more comprehensive direct comparison. Furthermore, these findings caution against using broad generalizations regarding the role of autophagy in T cell function.

Although it has been established that CD4+ T cells require high levels of mTORC1 activity and autophagy for optimal activation [8,9,19], the mechanisms that enable their parallel activation remain poorly understood. Here we demonstrated that CD3/CD28-stimulation induces autophagy, AMPK- and mTORC1-activity simultaneously during activation (**Figures 2A, 2D and S4B**), rewiring the canonical AMPK-ULK1/2-mTORC1 signaling axis [50]. mTORC1 was observed to function as expected in relation to the ULK1/2 complex during activation, negatively phosphorylating ULK1/2 (**Figure 2C**). When activated, mTORC1 tethers to the lysosomal membrane forming a signaling hub that impairs lysosomal degradation and promotes anabolism [51,52]. Given the observed increase in lysosomal degradation of autophagosomes (**Figure 2A**) and endosomes (**Figure 4F**) during CD4+ T cell activation, this suggests that some degree of compartmentalization could be happening to permit mTORC1 activity and lysosomal degradation to occur [53]. While CD3/CD28 stimulation also increased AMPK activity (**Figures 2D and S3B**), there was no AMPK-dependent phosphorylation of ULK1/2 (**Figure 2E**). This suggests that the two complexes do not interact following TCR/co-stimulation, potentially due to mTORC1 phosphorylation. TCR/co-stimulation-induced autophagy was also found to be ULK1/2-independent (**Figures 2G, S3D and S4B**), defining a novel non-canonical autophagy pathway that bypasses mTORC1-mediated inhibition and permits concurrent activation of autophagy and mTORC1 signaling. Additionally, autophagy was found to be ATG13-independent (**Figure 3B**), with *ATG13* deletion having no significant effect on the ability of CD4+ T cells to activate, proliferate and produce cytokines (**Figures 2I, 2J, 3C and 3D**). While this suggests the ULK1/2 complex is bypassed during activation, we cannot exclude a potential role for the remaining complex components ATG101 or FIP200. FBS starvation-induced autophagy remains dependent on ULK1/2 and the phosphorylation of ATG14 (**Figure 2F and 2G**). Notably however, this was also AMPK-independent in CD4+ T cells (**Figures 2D and S5B**). Furthermore, inhibition of mTORC1 did not induce autophagy in unstimulated cells **(Figure 2B**), suggesting a basal rewiring of the AMPK-ULK1/2-mTORC1 signaling axis. Both these phenomena were also reported in CD8+ T cells [14], indicating that, while autophagic responses to activation might differ between CD4+ and CD8+ T cells, both could still share similarities in regulation of stress-induced autophagy programs.

Previous studies have shown that inhibition of PIK3C3 can impair activation and lead to the establishment of an anergic phenotype [9,25]. Here we show that TCR/co-stimulation-induced autophagy is also PIK3C3-dependent (**Figures 2H and S4C**), with PIK3C3 inhibition significantly reducing cytokine production and proliferation in activated cells (**Figures 2I, 2J and S4C**). TCR/co-stimulation-induced autophagy does not require ULK1/2-dependent phosphorylation of ATG14 (**Figure 2F**) and can occur independently of Beclin 1 (**Figure 3E**), suggesting the formation of PIK3C3-C1 might not be necessary for PIK3C3-dependent regulation. While Beclin 1 is dispensable for activation-induced autophagic flux, it may exert non-canonical inhibitory roles on T cell effector function, potentially through its involvement in alternative vesicle trafficking pathways. PIK3C3 also functions within an alternate complex, PIK3C3-C2, consisting of UV radiation resistance-associated gene (UVRAG), Beclin 1, PIK3R4 and PIK3C3. In this conformation the complex regulates autophagosome maturation and endosomal trafficking [54]. While Beclin 1 is still a core unit of PIK3C3-C2, it could be possible that the remaining members are still functioning to regulate autophagy. Immuno-EM images revealed that TCR/co-stimulation-induced autophagy is morphologically distinct (**Figures 4A, 4B and 4C**) forming abundant small autophagic structures that frequently appeared as dense, internalized LC3+ content in multivesicular bodies (**Figure 4D and 4E**). While we could not distinguish whether these autophagic structures were accumulating in amphisomes or autolysosomes, we did determine that the effect of PIK3C3 inhibition was not due to inhibition of endolysosomal trafficking (**Figure 4F**), confirming that we are observing a distinct autophagy pathway. The formation of dense LC3+ autophagic structures suggests that TCR/co-stimulation could be inducing a selective form autophagy during activation (**Figure 4C)** [55,56]. This would support previous observations made by us and others which indicate that during activation autophagy could function to selectively target cargo to enable activation and regulate signaling pathways, such as in the case of PTPN1 and IL-7Rα [9,10].

While other ULK1/2-independent forms of non-canonical autophagy have been described [29,31,57–59], none match the phenotype or fit within the parameters defined here. Examples of Beclin 1-independent autophagy are rare, being implicated in apoptosis and hypoxia [60–62], potentially making our observations unique to CD4+ T cell activation. As such it is still unclear what the precise upstream components that regulate autophagy during activation are. Here we show that PMA/ION can be used as an alternative stimulation to induce the same autophagic response (**Figures S4B and S4C**). PMA potently activates protein kinase C (PKC) isoforms and their downstream signaling pathways [28], and ION is a Ca^2+^ ionophore that promotes calcium signaling [27], which together mimic TCR and CD28 signaling to induce T cell activation. While this indicates TCR engagement is not necessary, further investigation is needed as to how PMA/ION and TCR/CD28 signaling converge to activate autophagy [63,64]. Thus far we can rule out AMPK as a potential activator (**Figures 2E and 3A**), as well as any regulators that exclusively interact with the canonical autophagy pathway through the ULK1/2 complex [22] (**Figures 2F and 2G**).

Taken together, our findings identify a previously unrecognized non-canonical autophagy pathway in CD4+ T cells, that operates independently of mTORC1, AMPK and the ULK1/2 complex, yet remains PIK3C3-dependent. This mechanism enables autophagy to occur alongside anabolic signaling and provides a potential explanation for how activation of these antagonistic pathways is coordinated during activation. Importantly, this unique pathway provides the possibility of specific therapeutic targeting of autophagy to modulate pathogenic CD4+ T cell responses.

## Materials and Methods

### Blood collection and CD4+ T cell isolation

Human CD4+ T cells were isolated from peripheral blood (PB) collected either from anonymous healthy donors enrolled in the Minidonor Dienst Program at the UMC Utrecht after prior informed consent, or purchased buffy coats (Sanquin, NL). The age range of the donors is unknown, but it is estimated to be between 25 and 60 years old. Additionally, as gender is not disclosed, it is unclear whether the samples were obtained from male or female individuals. The study procedures were approved by the Institutional Review Board of the University Medical Centre Utrecht (METC, 11-499c), and performed according to the principles expressed in the Declaration of Helsinki. PBMCs were isolated from PB using Ficoll-Paque Plus (Sigman Aldrich) density gradient centrifugation. Primary CD4+ T cells were isolated from PBMCs using MagniSort Human CD4 T cell Enrichment Kit (Thermo Fisher Scientific), or CD4 Microbeads (Miltenyi) according to the manufacturer’s instructions.

### Cell culture, stimulations and inhibitors

CD4+ T cells were cultured in RPMI 1640 Medium with GlutaMAX™ (Thermo Fisher Scientific), supplemented with 10% FBS (Thermo Fisher Scientific) and 100 U/mL penicillin and 100 μg/mL streptomycin (Thermo Fisher Scientific). Cells were maintained at 37°C in a 5% CO2 humidified atmosphere. For CD3/CD28 activation, cells were stimulated with 5 ul/ml T Cell TransAct (Miltenyi), or 1 µg/ml anti-CD3 (eBioscience, 16-0037-81) and 1 µg/ml anti-CD28 (eBioscience, 16-0289-81). Where indicated, cells were activated with 20 ng/ml PMA (Sigma-Aldrich) and 1 µg/ml ION (Sigma-Aldrich). For activations cells were stimulated for 24 h unless otherwise stated. For cytokine treatments, cells were cultured with recombinant human IL-2 (PrepoTech), IL-7 (Miltenyi, premium grade), or IL-15 (Miltenyi, premium grade) at the concentrations indicated in the figures or figure legends. The following inhibitors were used for the pharmacological inhibition of Signaling pathways: Rapamycin (Sigma-Aldrich), Torin 1 (Sigma-Aldrich), 1 µM MRT68921 (Sigma-Aldrich), 1 µM ULK101 (Selleckchem), 1 µM Autophinib (Selleckchem), 0.5 µM PIKIII (Selleckchem), 2.5 µM LY294002 (Selleckchem), and 0.5 µM BAY-3827 (Selleckchem). Inhibitions were performed at the given concentrations unless otherwise stated. For starvation experiments, cells were cultured in RPMI 1640 Medium (Thermo Fisher Scientific), supplemented with 1% FBS (Thermo Fisher Scientific), and 100 U/mL penicillin and 100 μg/mL streptomycin (Thermo Fisher Scientific) for 24 h.

### Immunoblotting

CD4+ T cells were washed with ice-cold PBS and lysed in RIPA lysis buffer (150mM NaCl, 1% Triton X-100, 0.5% sodium deoxycholate, 0.1% SDS, 50mM Tris, pH 8.0) containing 1x Halt protease inhibitor and 1x Halt phosphatase inhibitor (Thermo Fisher Scientific). Protein concentration was determined using Bradford protein assay (Bio-Rad). Lysates were mixed with 1x western-ready protein sample loading buffer (Biolegend) with 2% β-mercaptoethanol and boiled at 100°C for 5 min. Equal amounts of sample (20-30 µg) were separated by SDS-PAGE on 8, 10, 12 or 15% acrylamide gels, and transferred to PVDF membranes using a Trans-Blot Turbo System (Bio-Rad). Membranes were blocked with either 5% milk protein in TBST (20mM Tris pH 8, 30mM NaCl, 0.2% Tween-20), or with 3% bovine serum albumin (BSA) (Sigma-Aldrich) in TBST for staining phospho-antibodies. Membranes were incubated 24 h at 4°C, in 3% BSA/TBST with primary antibodies LC3B (MBL, PM036, 1:1000), LC3B (NanoTools, 0231-100, 1:1000), S6 (54D2) (Cell Signaling, 2317, 1:1000), pS6 s235/236 (Cell Signaling, 2211, 1:1000), β-Actin (C4) (Santa Cruz, sc-47778, 1:1000), AMPKα (F6) (Cell Signaling, 2793, 1:1000), pAMPKα t172 (Cell Signaling, 2531, 1:1000), ULK1 (D8H5) (Cell Signaling, 8054, 1:500), pULK1 s757 (Cell Signaling, 6888, 1:500), pULK1 s555 (D1H4) (Cell Signaling, 5869, 1:500), ATG14 (Cell Signaling, 5504, 1:500), pATG14 s29 (Cell Signaling, 13155, 1:1000), ACC (C83B10) (Cell Signaling, 3676, 1:500), pACC s79 (D7D11) (Cell Signaling, 11818, 1:500), AKT (Cell Signaling, 9272, 1:1000), pAKT s473 (Cell Signaling, 9271, 1:1000), Vinculin (7F9) (Thermo Fisher Scientific, 14-9777-82, 1:5000), ATG13 (D4P1K) (Cell Signaling, 13273, 1:1000) or Beclin 1 (Cell Signaling, 3738, 1:1000). After washing with TBST, membranes were stained for 1 h at room temperature in 5% milk/TBST with secondary antibodies rabbit anti-mouse immunoglobulin-HRP (Dako, P0260, 1:5000), swine anti-rabbit immunoglobulin-HRP (Dako, P0217, 1:5000) or IRDye 680RD Donkey Anti-Mouse IgG (LICORbio, 926-68072, 1:20,000). Membranes were washed with TBST before detection. HRP-stained membranes were imaged either using SuperSignal West Pico PLUS Chemiluminescent Substrate (Thermo Fisher Scientific), SuperSignal West Dura Extended Duration Substrate (Thermo Fisher Scientific), SuperSignal West Femto Maximum Sensitivity Substrate (Thermo Fisher Scientific) or SuperSignal West Atto Ultimate Sensitivity Substrate (Thermo Fisher Scientific), with a ChemiDoc™ Touch Imaging System (Bio-Rad). Fluorescent-stained membranes were imaged using an Odyssey Imaging System (LICORbio). Images were processed using Image Lab (Bio-Rad) or Adobe Photoshop, and quantified in FIJI using integrated density.

### Flow Cytometry and Cell Sorting

To measure proliferation with flow cytometry, CD4+ T cells were activated with anti-CD3/CD28 for either 24 h or 48 h. They were then washed with PBS, and stained with live/dead marker Zombie NIR Fixable Viability Kit (Biolegend, 1:1000) in PBS for 15 min. Cells were subsequently washed in PBS, and intracellular Ki67 staining was performed using the Intracellular Fixation & Permeabilization Buffer Set (eBioscience) according to the manufacturers protocol, with Ki67 (Abcam, ab15580, 1:200) and Alexa Fluor 488 Donkey anti-rabbit IgG (Biolegend, 406416, 1:200). To stain for cell surface activation markers, cells were activated with anti-CD3/CD28 for 24 h, and then washed in PBS and stained with live/dead marker Zombie NIR Fixable Viability Kit (Biolegend, 1:1000) in PBS for 15 min. Cells were then washed in PBS, and incubated with anti-human CD25 FITC-conjugated (Immunotools, 21810253, 1:100) and Brilliant Violet 605 anti-human CD69 (Biolegend, 310938, 1:100) for 20 min in MACs buffer (2% FBS, 2mM EDTA, in PBS). Lysosomal activity was assessed using DQ Green BSA (Thermo Fisher Scientific). Cells were activated with anti-CD3/CD28 for 24 h, and cultured with 10 µg/mL DQBSA for 6 h. For all experiments, cells were washed in MACs buffer prior to acquisition. Results were acquired on a BD LSRFortessa Cell Analyzer (BD Biosciences) with FACSDiva (BD Biosciences) software or a CytoFLEX Flow Cytometer (Beckman Coulter) with CytExpert software. Data was analysed with FlowJo (BD Biosciences).

For sorting naïve and memory T cell subsets, CD4+ T cells were stained with APC anti-human CD45RA (Biolegend, 304150, 1:200) and FITC anti-human CD45RO (Biolegend, 304204, 1:200) for 20 min in MACs buffer. Cells were washed in MACs buffer and sorted using a BD Influx Cell Sorter (BD Biosciences). Restrictive gating was applied to ensure only single positive cells were sorted in each channel. Cells were rested in culture 24 h after sorting prior to experimental use.

### Autophagic flux and saponin extraction

To assess autophagic flux, CD4+ T cells were treated with ± 100 nM BafA1(Sigma-Aldrich) for the last 3 h of culture prior to analysis. Autophagic flux was either measured using western blot, or flow cytometry with saponin extraction. For western blot, cells were lysed and immunoblotted as described above, using high percentage 12-15% acrylamide gels to ensure the separation of LC3II [20]. For flow cytometry, saponin extraction was performed to flush non-autophagosomal LC3 as previously described [21]. Briefly, cells were washed with PBS, followed by saponin buffer (0.1% saponin [Sigma-Aldrich] in PBS). They were then stained with LC3B (MBL, PM036, 1:200) in saponin buffer for 20 min. Cells were washed in saponin buffer, and then stained with Alexa Fluor 647 Donkey anti-rabbit IgG (Biolegend, 406414, 1:100) in saponin buffer for 20 min. Cells were washed in MACs buffer, and acquired on a BD LSRFortessa Cell Analyzer (BD Biosciences) with FACSDiva (BD Biosciences) software. Data was analysed with FlowJo (BD Biosciences).

### CRISPR-Cas9 CD4+ T cell genome editing

Predesigned sequences of Alt-R crRNA specific to BECN1 and ATG13 were purchased from IDT. Sequences as follows: *BECN1:* ATCTGCGAGAGACACCATCC, on-target score: 58, off-target score: 68. *ATG13* :CTGTCCCAACACGAACTGTC, on-target score: 65, off-target score: 84. For preparation of RNPs, crRNA were resuspended at 160 µM in IDT RNA duplex buffer and mixed in a 1:1 ratio with 160 µM Alt-R tracrRNA (IDT, 1072533). The mix was incubated at 95°C and allowed to cool down to room temperature, to generate annealed gRNA at 80 µM concentration. To improve RNP formulation, a solution of single-stranded oligodeoxynucleotide (ssODN) was added as previously described [65]. This molecule was ordered as a single-stranded Ultramer from IDT (sequence: 5’ TTAGCTCTGTTTACGTCCCAGCGGGCATGAGAGTAACAAGAGGGTGTGGTAATATTACGGTACCGAGCACTATCGATACAATATGTGTCATACGGACACG 3’) and resuspended at 100 µM in nuclease-free water. gRNA and ssODN were mixed in equimolar amounts (1:0.8 volume ratio). Cas9 protein (QB3 Macrolab, University of California Berkeley) was added to the gRNA in a 1:2 ratio and incubated at room temperature for 15 min to generate RNPs. RNPs were used either after formulation or stored at -20 °C and thawed once for repeated use.

For CRISPR KO experiments, CD4+ T cells were activated with TransAct in TexMACS culture medium (Miltenyi) with 2% human serum and 100 U/mL penicillin and 100 μg/mL streptomycin (Thermo Fisher Scientific) with 5 ng/ml of IL-7 (Miltenyi, premium grade) and 5 ng/ml IL-15 (Miltenyi, premium grade). After two days of activation, cells were washed with PBS and resuspended in supplemented P3 buffer (Lonza) at a concentration of 50×10e^6^/ml. RNP were added to a final concentration of 2 µM and 20 µl of cells were electroporated in a 16-well strip. Following electroporation, cells were rested in TexMACS medium with 5% human serum and no antibiotics for 1-4 hours after which time pen/strep was added together with IL-7 and IL-15 to 5ng/ml. Cells were expanded for 4-6 days before analysis.

### ELISA

IL-2 production was measured using a ELISA MAX Deluxe Set Human IL-2 kit (Biolegend) according to the manufacturer’s instructions. Culture medium was harvested 24 h after activation. Standards, controls and samples were assayed in technical triplicates. The optical density of the color was measured with a microplate reader set at 450 nm. Cell proliferation was assessed by BrdU incorporation using a Colorimetric Cell Proliferation ELISA (Roche) according to the manufacturer’s protocol. Samples were assayed in technical triplicates. Cells were activated and incubated with BrdU reagent for 24 h prior to harvest and assay. Absorbance was measured at 450 nm using a microplate reader after the addition of a stop solution (0.2 M H_2_SO_4_).

### Immuno-electron microscopy

CD4+ T cells were treated with ± 20 nM BafA1 (Sigma-Aldrich) for the last 2 h prior to fixation and analysis. They were fixed in 4% paraformaldehyde and processed for ultrathin cryosectioning as previously described [66]. Sections were immunolabeled using LC3B clone LC3-1703 (Cosmo Bio, CAC-CTB-LC3-2-IC), rabbit anti-mouse IgG (Rockland, 610–4120, 1:250) and protein A-10 nm gold particles made in house (Cell Microscopy Core, UMC Utrecht). Images were acquired using a Tecnai T12 (FEI Tecnai) and SerialEM software (Nexperion). The images were further processed in ImageJ. Quantification was performed on compartments with at least 2 gold particles. Cell profiles were screened for these compartments, and, if present, their surface area and amount of gold particles were quantified.

### Statistical analysis

Data are presented as mean ± SEM. Statistical significance was determined using one-way ANOVA with Tukey’s post-hoc test for multiple comparisons or Welch’s t-test for pairwise comparisons, as indicated in the figure legends. *p* < 0.05 was considered statistically significant (\**p* < 0.05, \*\**p* < 0.01, \*\*\**p* < 0.001, \*\*\*\**p* < 0.0001). All analyses were performed using GraphPad Prism 9 software.

## Acknowledgements

We would like to thank members of the Coffer Lab for valuable discussions and Suzanne van Dijk and Cecilia de Heus, for technical assistance. This work was supported by a grant from the ReumaNederland (18-1-401).

## Supplementary Figures

**Supplementary Figure S1.**
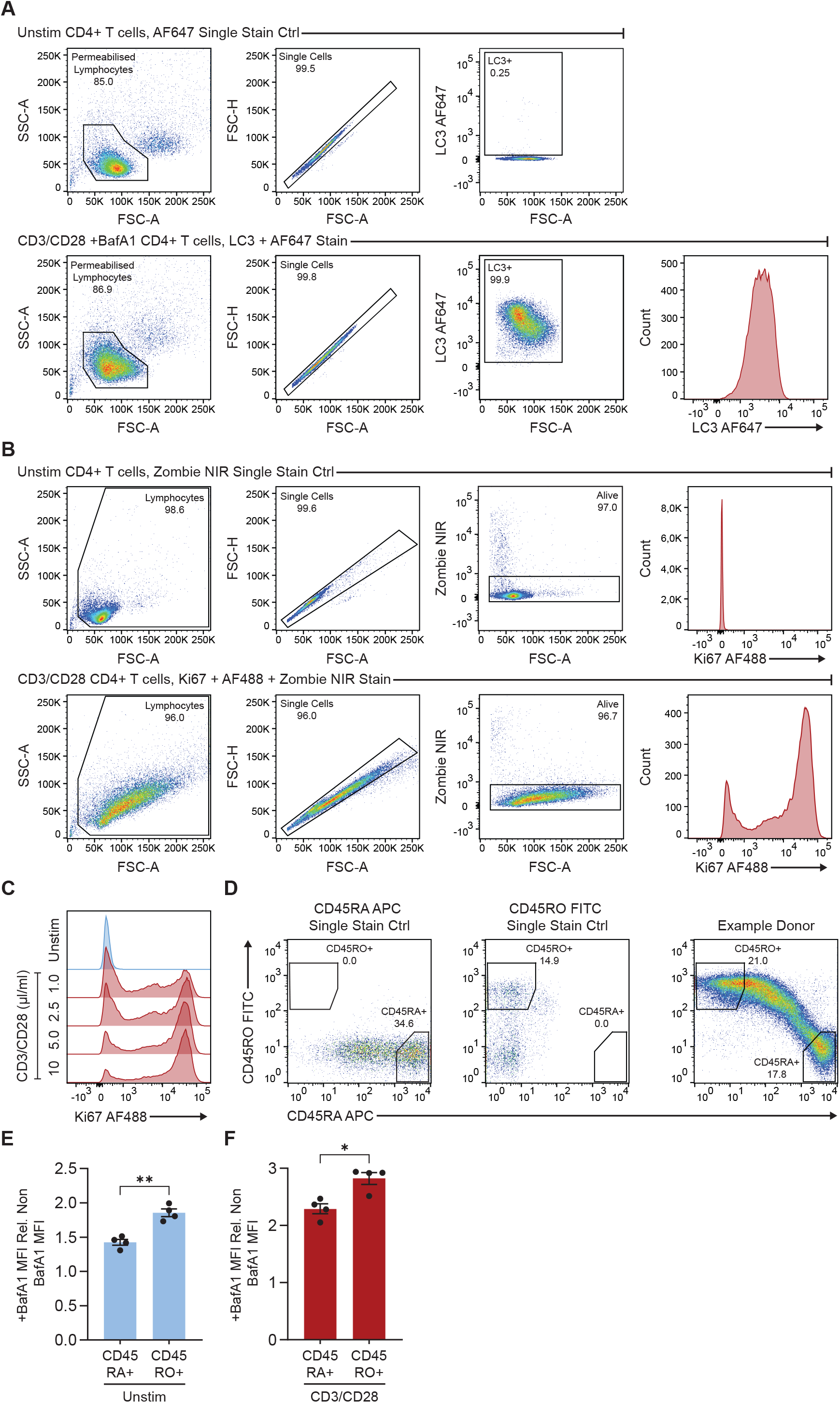
(**A**) Representative gating strategy for autophagic flux saponin assay. CD4+ T cells were gated for permeabilized lymphocytes, single cells and LC3+ cells before being displayed as normalized modal histograms. Gates were set based on single stain controls. Examples shown are CD4+ T cells either, unstimulated 24 h and stained only with AF647 secondary antibody, or stimulated 24 h with anti-CD3/CD28, treated with 100 nM BafA1 for 3 h and stained with LC3 primary and AF647 secondary antibodies. (**B**) Representative gating strategy for Ki67 proliferation assay, showing size and live/dead gating. CD4+ T cells were gated for lymphocytes, single cells and alive cells before being displayed as normalized modal histograms. Gates were set based on single stain controls. Examples shown are for CD4+ T cells either, unstimulated 48 h and stained only with Zombie NIR live/dead stain or stimulated 48 h with anti-CD3/CD28 and stained with Zombie NIR live/dead stain, Ki67 primary and AF488 secondary antibodies. (**C**) Representative normalized modal histograms of Ki67 MFI for Figure 1E. (**D** and **E**) Relative level of autophagic flux as measured through flow cytometry using saponin extraction and LC3II MFI and determined by the ratio of LC3II in the +BafA1 condition and non BafA1 condition between CD45RA+ and CD45RO+ sorted CD4+ T cells. Cells were either (**E**) Unstimulated 24 h, or (**F**) Stimulated 24 h with anti-CD3/CD28, and treated 3 h ± 100 nM BafA1. Representative gating strategy for CD45RA+ and CD45RO+ sorting of CD4+ T cells. Cells were first gated based on size and single cells as in (**B**), before being sorted for CD45RA or CD45RO. Gates were set based on single stain controls. Examples shown are for CD4+ T cells either single stained only with CD45RA APC antibody, CD45RO FITC antibody or an example donor stained with CD45RA APC and CD45RO FITC antibodies. Gating was set restrictively to ensure only single positive cells in each condition were sorted. All graphs represent mean ± SEM. Statistical significance was measured by Welch’s t-test. ∗*p* < 0.05, ∗∗*p* < 0.01.

**Supplementary Figure S2.**
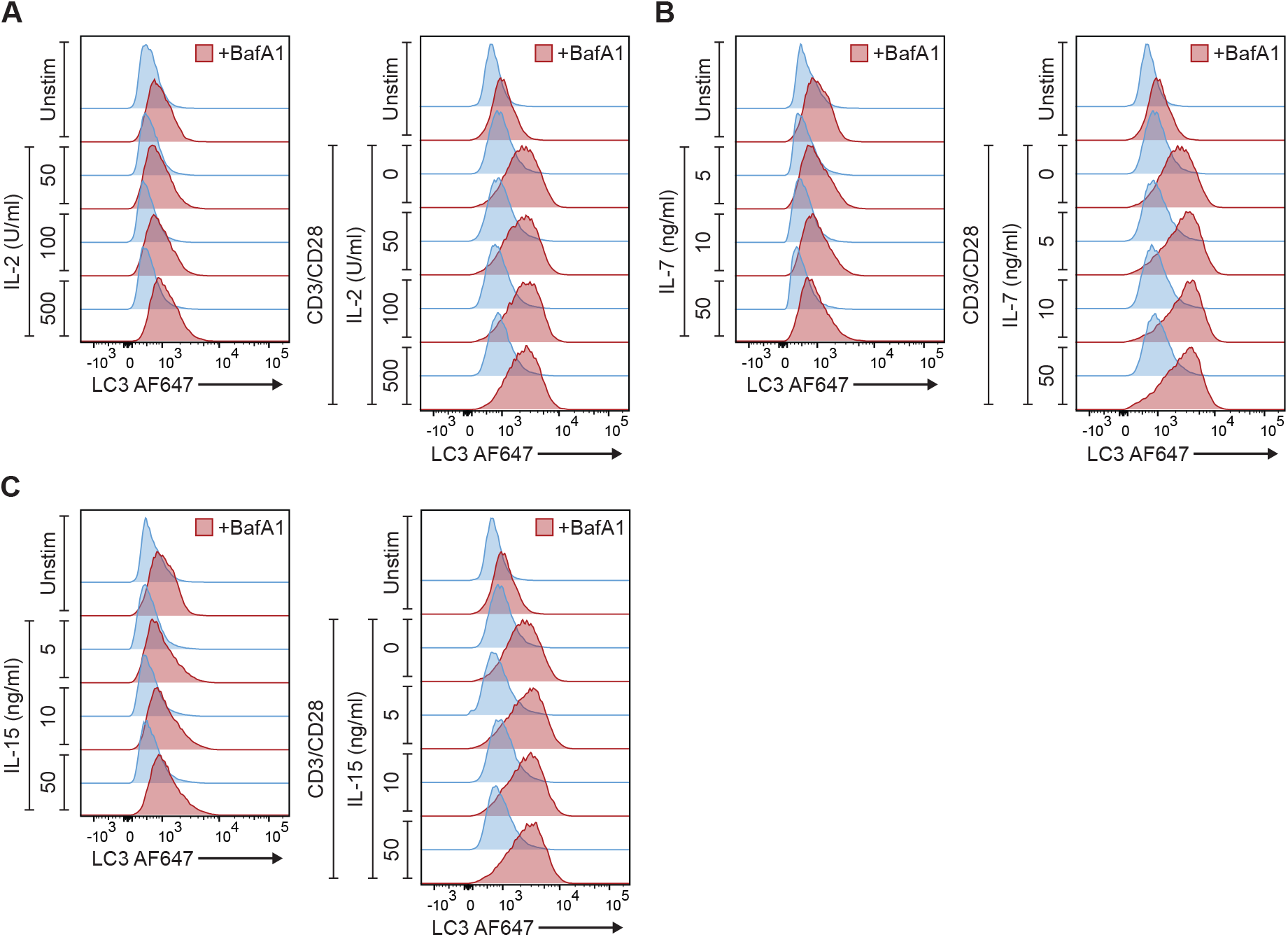
(**A-C**) Representative normalized modal histograms of autophagic flux, measured through flow cytometry using saponin extraction and LC3II MFI, for (**A**) Figure 1H, Figure 1I and (**C**) Figure 1J.

**Supplementary Figure S3.**
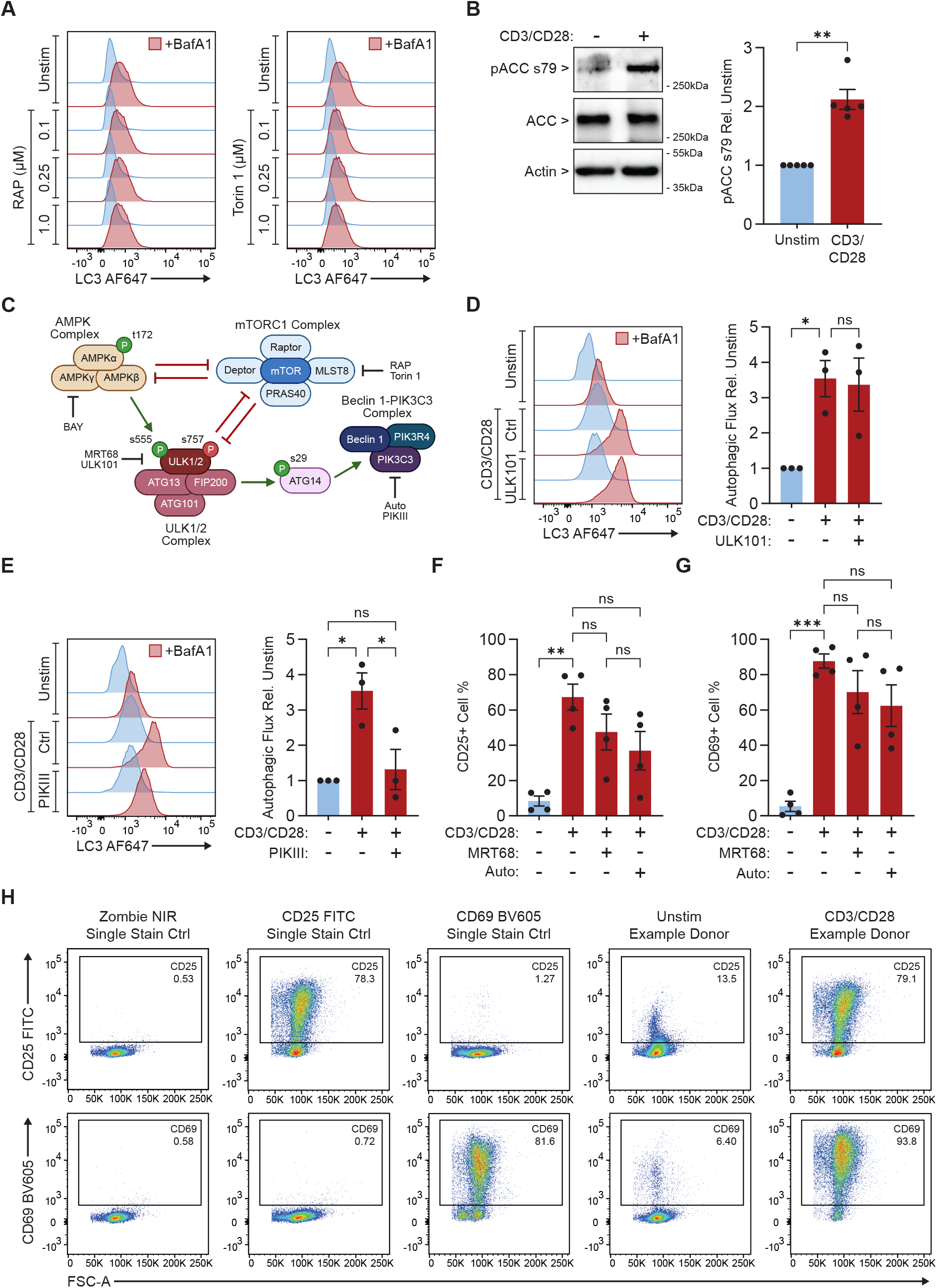
(**A**) Representative normalized modal histograms of autophagic flux, measured through flow cytometry using saponin extraction and LC3II MF, for figure 2B. (**B**) CD4+ T cells stimulated 24 h with anti-CD3/CD28. Representative western blot of pACC s79, with quantifications (n=5). (**C**) Schematic overview of the autophagy pathway interactions examined in this study, and the targets of the inhibitors used. The AMPK αβγ heterotrimer complex is composed of the catalytic α subunit, and the regulatory and scaffolding β and γ subunits. Phosphorylation of AMPK t172 is a key activation marker of complex activity. AMPK negatively regulates the mTORC1 complex and positively regulates ULK1/2 activity through the phosphorylation of ULK1/2 s555. mTORC1 is comprised of mTOR, regulatory-associated protein of mTOR (Raptor), DEP domain-containing mTOR-interacting protein (Deptor), target of rapamycin complex subunit LST8 (MLST8) and proline-rich AKT substrate of 40 kDa (PRAS40). mTORC1 negatively regulates AMPK activity, and ULK1/2 through the phosphorylation of ULK1/2 s757. The ULK1/2 complex is made up of ULK1/2, ATG13, FIP200 and ATG101. It negatively regulates mTORC1 and promotes autophagy through the phosphorylation of ATG14 s29 to assemble the PIK3C3-C1 complex with Beclin 1, PIK3R4 and PIK3C3. (**D and E**) Representative normalized modal histograms of autophagic flux measured through flow cytometry using saponin extraction and LC3II MFI, with quantifications (n=3), of CD4+ T cells stimulated 24 h with anti-CD3/CD28 and either treated with (**D**) 1 µM ULK101 or (**E**) 0.5 µM PIKIII, and ± 100 nM BafA1 for 3 h. (**F and G**) CD4+ T cells stimulated 24 h with anti-CD3/CD28 and treated with 1 µM Auto or 1 µM MRT68. Graphs show: (**F**) Percentage of CD25+ cells measured through flow cytometry; (**G**) Percentage of CD69+ cells measured through flow cytometry. (**H**) Representative gating strategy for CD25 and CD69 cell surface marker expression. CD4+ T cells were first gated for size, single cells and alive cells as in Figure S1B. Gates were set based on single stain controls. Examples shown display single stain controls or example donors. CD4+ T cells were either unstimulated or stimulated 24 h with anti-CD3/CD28, and single stained with Zombie NIR live/dead, CD25 FITC antibody or CD69 BV605 antibody, or stained with Zombie NIR live/dead, CD25 FITC and CD69 BV605 antibodies. All graphs represent mean ± SEM. Statistical significance was measured by one-way ANOVA with Tukey post-hoc test. ∗*p* < 0.05, ∗∗*p* < 0.01, ∗∗∗*p* < 0.001.

**Supplementary Figure S4.**
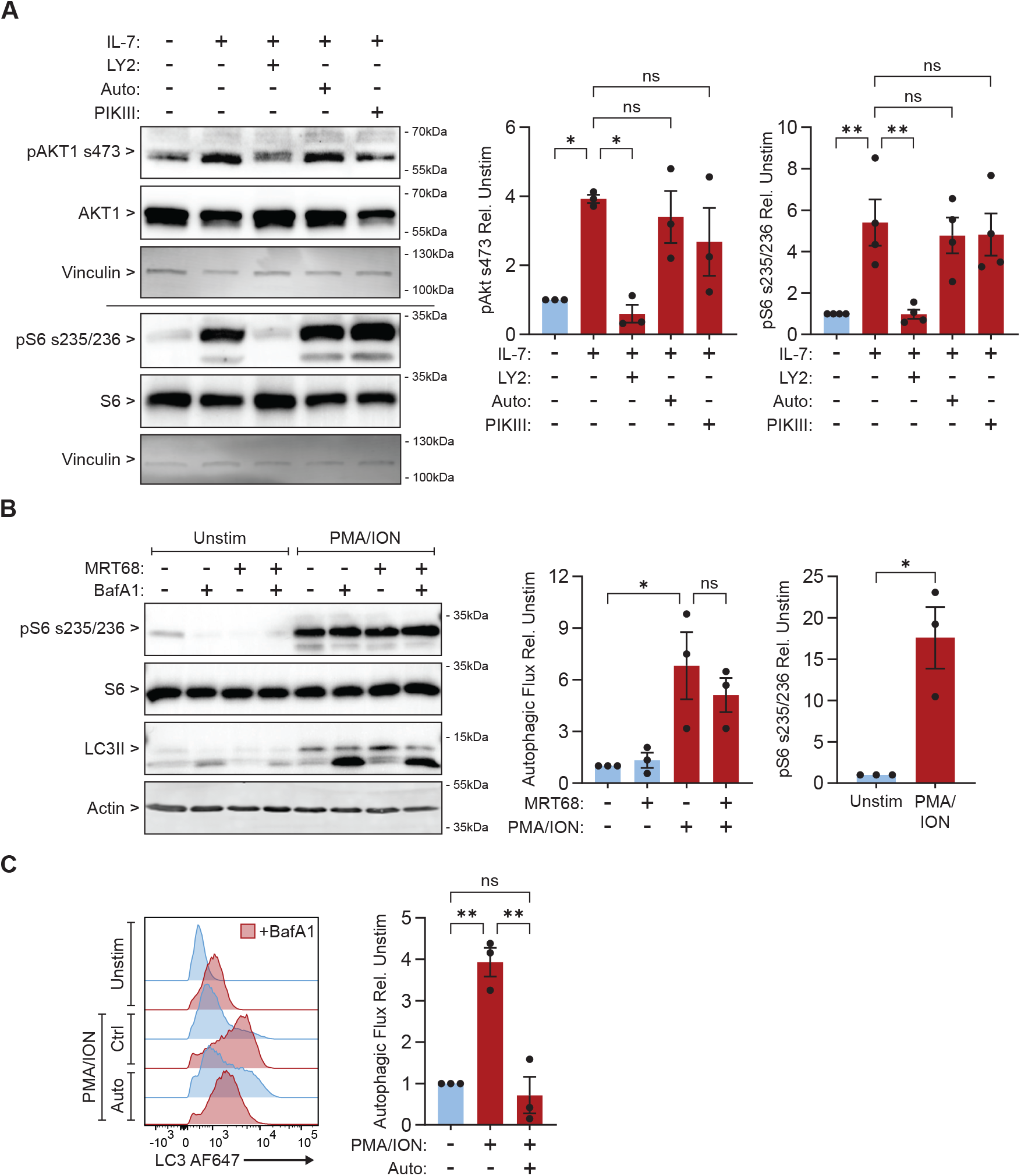
(**A**) CD4+ T cells stimulated 4 h with 5 ng/ml IL-7 and treated either with 2.5 µM LY294002 (LY2), 1 µM Auto or 0.5 µM PIKIII. Representative western blots with quantifications of downstream PI3K activity as measured by levels of pAKT1 s473 (n=3), and pS6 s235/236 (n=4). (**B**) CD4+ T cells stimulated 24 h with PMA/ION and treated with ± 1 µM MRT68 for 24 h, and ± 100 nM BafA1 for 3 h. Representative western blot and quantifications (n=3) of autophagic flux and mTORC1 activity, as measured by LC3II turnover and pS6 s235/236 levels. Representative normalized modal histograms of autophagic flux measured through flow cytometry using saponin extraction and LC3II MFI, with quantifications (n=3), of CD4+ T cells stimulated 24 h with PMA/ION and treated with 1 µM Auto and ± 100 nM BafA1 for 3h. All graphs represent mean ± SEM. Statistical significance was measured by one-way ANOVA with Tukey post-hoc test. ∗*p* < 0.05, ∗∗*p* < 0.01.

**Supplementary Figure S5.**
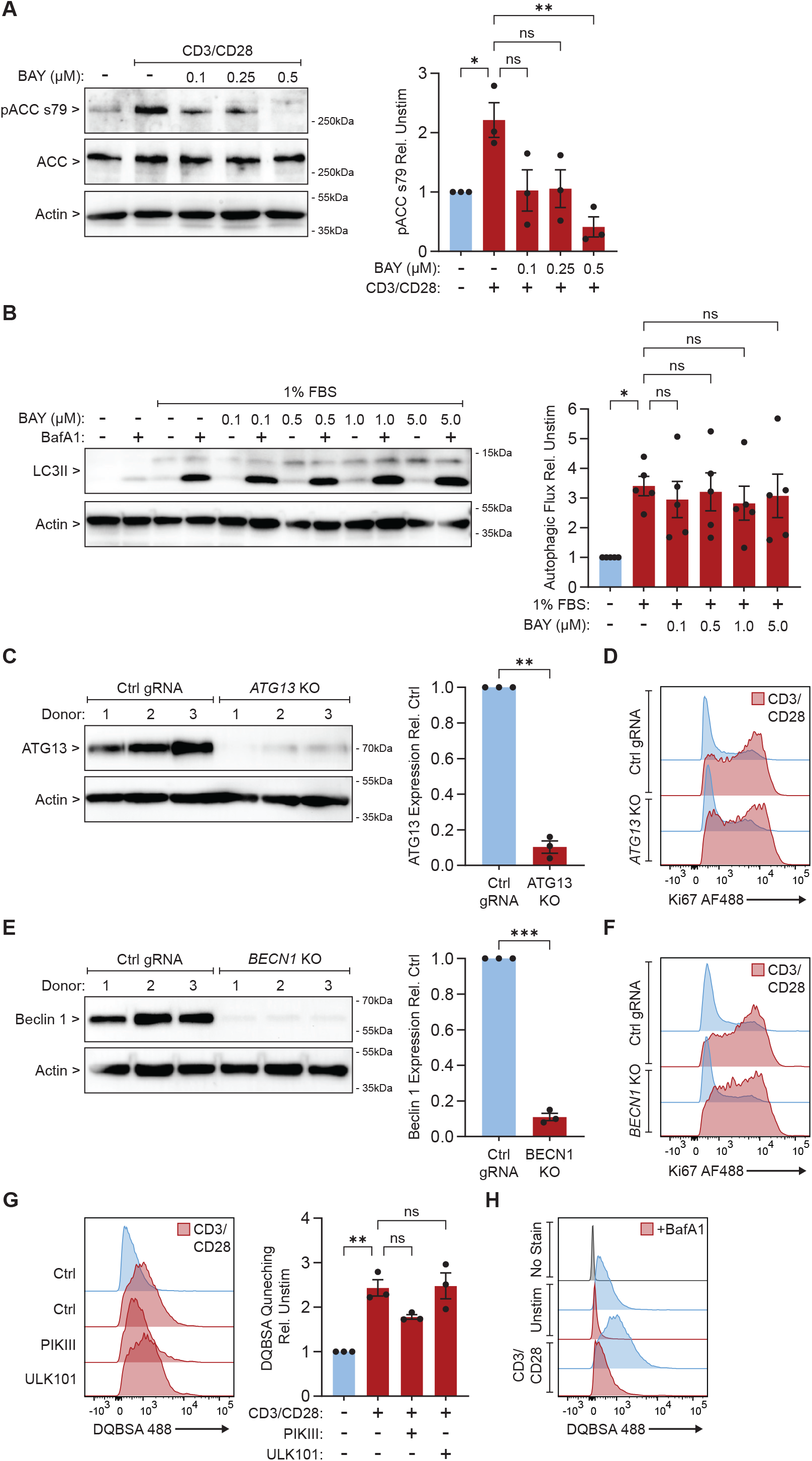
(**A**) CD4+ T cells stimulated 24 h with anti-CD3/CD28 and treated with increasing concentrations of BAY. Representative western blot of pACC s79 levels, with quantifications (n=3). (**B**) CD4+ T cells starved with 1% FBS and treated with different concentrations of for 24 h and ± 100 nM BafA1 for 3 h. Representative western blot of autophagic flux as measured by LC3II turnover, with quantifications (n=5). (**C**) Western blot for ATG13 KO efficiency in CD4+ T cells across 3 donors, with quantifications. (**D**) Representative normalized modal histograms of Ki67 MFI for Figure 3E. (**E**) Western blot for BECN1 KO efficiency in CD4+ T cells across 3 donors, with quantifications. (**F**) Representative normalized modal histograms of Ki67 MFI for Figure 3H. (**G**) CD4+ T cells stimulated 24 h with anti-CD3/CD28 and treated with 0.5 µM PIKIII or 1 µM ULK101, and cultured 6 h with 10 µg/ml DQBSA. Representative normalized modal histograms of DQBSA quenching, with DQBSA MFI quantifications (n=3). (**H**) CD4+ T cells stimulated with anti-CD3/CD28 and treated ± 100 nM BafA1 for 24 h, and cultured 6 h with or without 10 µg/ml DQBSA. Normalized modal histograms of DQBSA dequenching (No Stain = without DQBSA). All graphs represent mean ± SEM. Statistical significance was measured by one-way ANOVA with Tukey post-hoc test, or by Welch’s t-test. ∗*p* < 0.05, ∗∗*p* < 0.01, ∗∗∗*p* < 0.001.

## Notes

### Competing Interest Statement

The authors have declared no competing interest.

